# Hydrogen sulfide increases intracellular oxygen and regulates the HIF response

**DOI:** 10.1101/2025.07.24.666576

**Authors:** Joseph Brake, David A Hanna, Roshan Kumar, Qianni Peng, Aaron P. Landry, Rashi Singhal, Eranthie Weerapana, Yatrik M. Shah, Ruma Banerjee

## Abstract

O_2_ sensing by hypoxia-inducible factor (HIF) is a principal mechanism by which aerobic organisms adjust cellular energy metabolism and adapt to O_2_ limitation. In this study, we show that H_2_S, a product of host and microbial metabolism, profoundly influences the threshold for HIF-dependent hypoxia-sensing by increasing intracellular O_2_. The dose-dependent destabilization of HIF by H_2_S is inversely correlated with sulfide quinone oxidoreductase, which oxidizes sulfide in the mitochondrion. Hypoxia sensors provide a quantitative estimate of the magnitude of H_2_S-induced perturbation. The O_2_ concentration in cells grown in a 2% O_2_ atmosphere is sensed as 5 or 15 % O_2_ in the presence of 25 or 100 ppm H_2_S, respectively. Sustained exposure to H_2_S elicits the hallmarks of hyperoxia-associated cytotoxicity, including loss of Fe-S proteins in cellular and murine models. H_2_S thus emerges as a powerful regulator of O_2_ sensing and signaling with possible implications for dysregulation in O_2_ toxicity diseases.

**Significance Statement:** The mitochondrial electron transport chain (ETC) accounts for ∼90% of whole body O_2_ consumption. However, our understanding of how metabolites modify ETC flux and therefore, intracellular O_2_ availability, are poor. In this study, we demonstrate that hydrogen sulfide (H_2_S), which is produced by host and gut microbes alike, increases intracellular O_2_ by decreasing ETC flux, and destabilizes the principal hypoxia sensor, HIF-1α. The upshift in intracellular O_2_ levels is quantitatively significant, such that 2% O_2_ is sensed as 5-15% O_2_ at varying H_2_S concentrations, with concomitant destabilization of Fe-S proteins, a signature of cellular hyperoxia. Our study identifies H_2_S as a HIF-1α regulator with important implications for the large class of mitochondrial diseases characterized by dysregulated O_2_ metabolism.

## INTRODUCTION

Biospheric oxygenation, which occurred ∼2.2 billion years ago, introduced a high-potential electron acceptor and substrate, fueling the evolution of new biocatalytic reactions and pathways. Today, O_2_ is estimated to be the most used substrate across metabolomes, surpassing even ATP and NADH (1). Compared to 21% O_2_ (160 mm Hg) in the atmosphere, O_2_ levels in human tissue vary from ∼14% O_2_ (110 mm Hg) in lung alveoli to <0.1% O_2_ (<1 mm Hg) in the intestinal lumen (2–4). Hypoxia inducible factor (HIF) is a principal O_2_ sensor, activating an adaptive transcriptional program for survival in low O_2_ (5–7). Under normoxic conditions, O_2_-dependent hydroxylation of the α-subunit of HIF by prolyl hydroxylase (PHD) tags it for ubiquitination by the von Hippel-Lindau (VHL) tumor-suppressor protein, a component of an E3 ligase complex, for subsequent proteasomal degradation (8, 9). An estimated 90% of whole body O_2_ consumption occurs at the electron transport chain (ETC) (10). Flux modulation and maintenance of tissue O_2_ gradients by endogenous ETC regulators are poorly understood. A mouse model for Leigh syndrome deficient in the complex I subunit Ndufs4 shows evidence of brain hyperoxia that can be reversed by hypoxia (11), highlighting the importance of ETC activity in O_2_ homeostasis.

The motivation of the current study was to investigate how H_2_S, which is a product of our own metabolism (12) and is also produced in copious quantities by gut microbes (13, 14), impacts O_2_ metabolism. In addition to being a respiratory poison that targets cytochrome c oxidase or complex IV (15), H_2_S is also an ETC substrate (16). In colon, the O_2_ gradient is shaped by the complex interaction between host and microbial metabolism and colonocytes occupy the liminal zone between a microbiota-dense and virtually anoxic lumen and a highly vascular lamina propia (17). The microbiome responds to luminal O_2_ levels and the O_2_ gradient in turn, affects host-microbiome interactions (2, 18). Chronic pathological hypoxia is a signature of active inflammatory bowel disease and if unchecked, contributes to disease progression (19). H_2_S is reported to both attenuate and augment HIF stability (20–23). Like nitric oxide, H_2_S reportedly induces metabolic hypoxia and redistributes O_2_ usage (20, 24).

In this study, we demonstrate that H_2_S elicits a dose-dependent destabilization of HIF under hypoxia (2% O_2_) by increasing intracellular O_2_. The magnitude of HIF destabilization is inversely correlated with expression of the H_2_S-oxidizing enzyme, sulfide quinone oxidoreductase (SQOR), and exacerbated by its knockdown. By calibrating intracellular O_2_ exposure in cells grown under ambient hypoxia and varying levels of H_2_S, we provide quantitative insights into the modulation of the threshold for hypoxia sensing. Further, while sustained H_2_S-dependent complex IV inhibition induces a reductive shift in redox cofactor pools (25), we report a concomitant oxidative shift in lipids as well as in the mitochondrial cysteine proteome accompanied by decreased levels of iron-sulfur (Fe-S) cluster proteins, a fingerprint of hyperoxia. Our study reveals that H_2_S is a powerful regulator of intracellular O_2_ availability, O_2_ sensing and HIF signaling.

## RESULTS

### Hypoxia restores Fe-S cluster proteins destabilized by H_2_S

Decreased forward electron flow due to complex IV inhibition has pleiotropic effects, including a reductive shift in redox cofactor pools (25, 26) and potentially, enhanced ROS and reactive sulfur species generation due to increased electron leakage (Fig. 1A). We used the proximity labeling oxidative isotope-coded affinity tag (PL OxICAT) platform (27) to evaluate how H_2_S affects the cysteine thiol status in the mitochondrial matrix proteome. Relative to cells grown in the absence of H_2_S, a broad oxidative shift was observed in the presence of H_2_S (100 ppm, 24 h, corresponding to ∼20 µM dissolved sulfide in the culture medium) (Fig. 1B, Supplementary Table 1). Notably, some of the oxidatively shifted targets were Fe-S proteins, including cluster-coordinating cysteine residues in NDUFS8 and SDHB, which are subunits of complex I and II, respectively (Fig. 1C). In addition, increased lipid peroxidation as reported by the BODIPY dye was observed in colon adenocarcinoma HT-29 cells exposed to H_2_S (Supplementary Fig. 1A,B). The magnitude of lipid oxidation elicited by H_2_S (100 ppm, 24 h) was comparable to that induced by cumene hydroperoxide (100 µM), employed as a positive control (Supplementary Fig. 1C,D).

**Figure 1:**
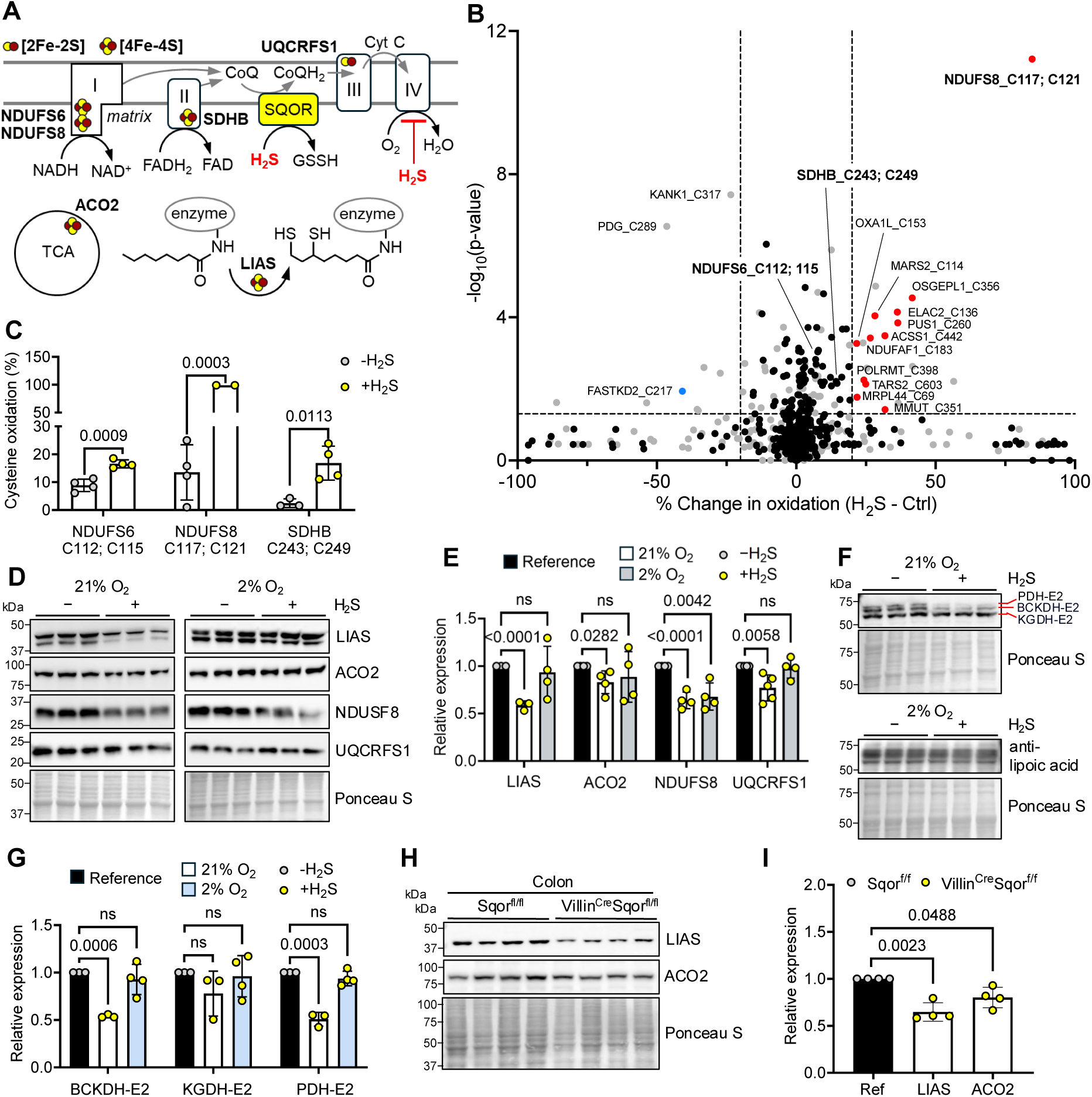
Hypoxia restores Fe-S proteins destabilized by H_2_S. (A) Model showing metabolic changes due to complex IV inhibition by H_2_S-mediated increase in O_2_, which can be mitigated by SQOR activity. Six Fe-S proteins that were found to be destabilized in this study by H_2_S are noted. (B) Volcano plot of PL-OxICAT data displaying the % change in oxidation for cysteines in HEK293 cells grown in the absence or presence of H_2_S (100 ppm, 24 h). The black and grey dots indicate mitochondrial and non-mitochondrial cysteines, and the red and blue dots indicate oxidized and reduced mitochondrial cysteines, respectively. Horizontal and vertical dashed lines indicate p value <0.05 and change in oxidation of 20%, respectively. (C) Quantitation of select Fe-S cluster-coordinating cysteines from the data in B. (D,E) Representative western blots (D) and quantitation (E) of select Fe-S proteins in HT-29 cells cultured at 21 or 2% O_2_ ± 100 ppm H_2_S (n=3 or 4). Reference denotes protein levels in the absence of H_2_S, arbitrarily set at 1. (F,G) Representative western blots (F) and quantitation (G) of lipoylated proteins in the same samples as D. (H,I) Representative western blots (H) and quantitation (I) of colon Fe-S proteins from Sqor^fl/fl^ (n=4) and *Villin*^Cre^ Sqor^fl/fl^ mice (n=4).

Next, we assessed the stability of mitochondrial Fe-S proteins with roles in lipid metabolism, the ETC and TCA cycle (Fig. 1A). A significant decrease in the expression of LIAS, ACO2, NDUFS8, and UQCRFS1 was seen in HT-29 cells grown in an atmosphere of 21% O_2_ and 100 ppm H_2_S (Fig. 1D,E). LIAS catalyzes the synthesis of lipoate, an essential cofactor for the E2 subunit of BCKDH, KGDH-E2 and PDH-E2, which are critical for branched chain amino acid catabolism and the TCA cycle. Lipoylated BCKDH and PDH subunits were decreased by H_2_S exposure in cells grown under normoxia (Fig 1F,G). Remarkably, with the exception of NDUFS8, hypoxia (2% O_2_) protected against H_2_S-triggered depletion of Fe-S proteins and LIAS targets (Fig. 1E,G), suggesting increased synthesis and/or stabilization of metal clusters under these conditions. The physiological relevance of Fe-S protein destabilization by elevated H_2_S exposure was assessed in *Villin*^Cre^ Sqor^fl/fl^ mice, which lacking the capacity for SQOR-dependent sulfide oxidation in intestinal epithelial cells, is expected to result in sustained exposure to higher levels of H_2_S(28) (Fig. 1A). LIAS and ACO2 levels were significantly lower in colon of *Villin*^Cre^ Sqor^fl/fl^ mice (Fig. 1H,I), consistent with observations in cell lines, although UQCRFS1 was unchanged (Supplemental Fig. 2A,B).

### H_2_S destabilizes HIF-1α under hypoxia

Restoration of Fe-S protein levels by shifting to a hypoxic atmosphere is consistent with our model that cluster damage results from sulfide-induced O_2_ accumulation (Fig. 1A). We therefore characterized the effect of H_2_S on hypoxia sensing. Acute exposure to Na_2_S (0.03-1 mM, 1 h) induced a dose-dependent decrease in HIF-1α stabilization in cells grown at 2% O_2_ (Fig. 2A,B). Since H_2_S disappears with a t^1^/_2_ of 6 min at 37 °C(29), the difference in HIF-1α between ± H_2_S samples was not seen beyond 1 h, which also revealed reversibility of the H_2_S effect (Supplementary Fig. 2C,D). Although HIF-1α has a molecular mass of 92.6 kDa, two major bands with estimated masses of 120 and 130 kDa were often, but not always, detected by Western blot analysis. Under hypoxic conditions, the less intense lower band was stabilized earlier and the difference was clearly seen in the high-exposure display (Supplementary Fig. 2C). H_2_S also destabilized HIF-1α in EA.hy296, HCT116 and HEK293 cells (Supplementary Fig. 2E-J).

**Figure 2:**
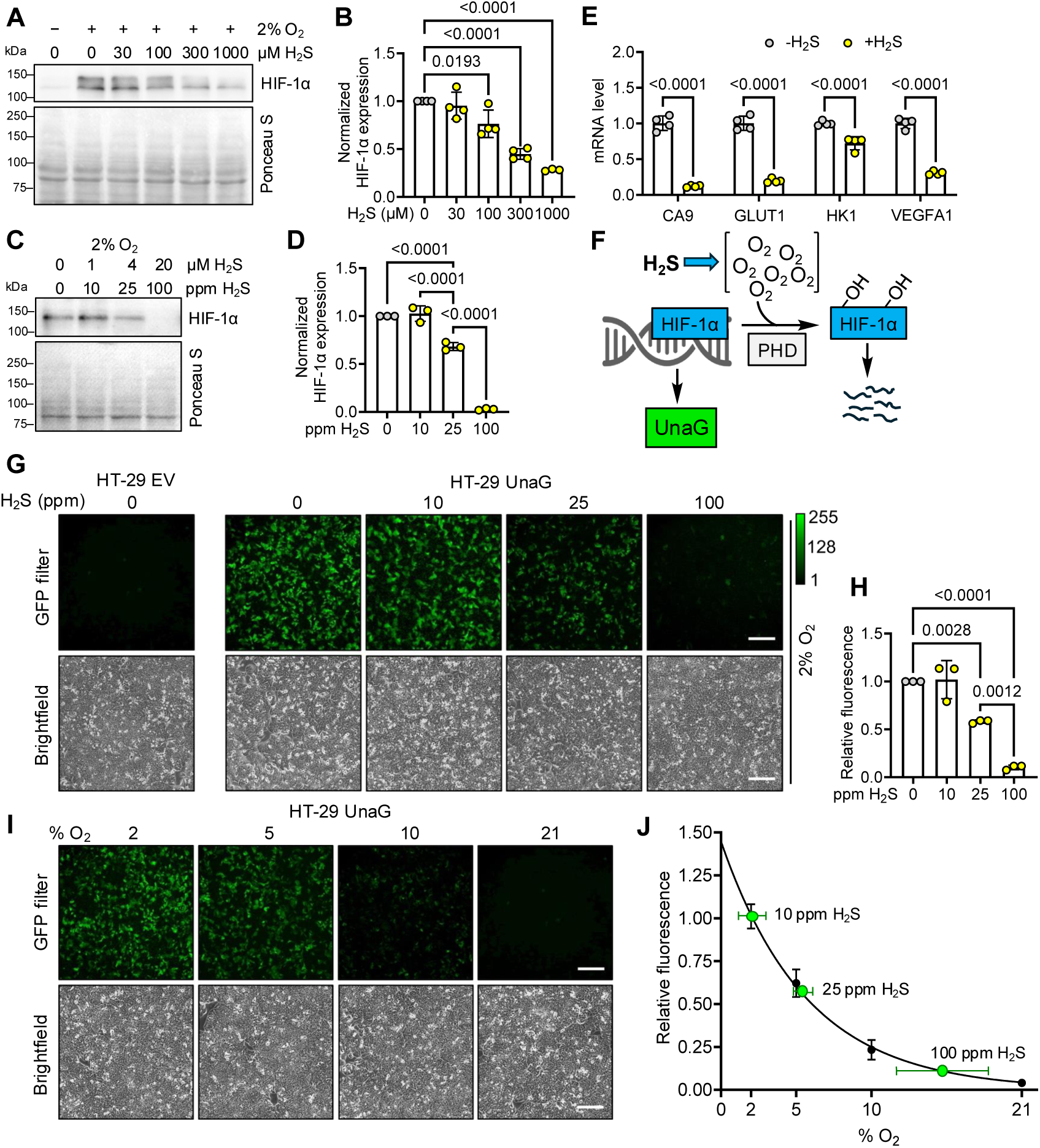
H_2_S shifts the cellular threshold for hypoxia sensing. (A) Representative western blot of HIF-1α in HT-29 cells grown under hypoxia (2% O_2_) with acute exposure to the indicated H_2_S concentration for 1 h (n=3 or 4). (B) Quantitation of data in A normalized to Ponceau S staining for total protein per lane. (C) Representative western blot of HIF-1α in HT-29 cells grown under hypoxia (2% O_2_) with chronic exposure to the indicated H_2_S concentration for 24 h. (D) Quantitation of data in C normalized to Ponceau S staining for total protein per lane (n=3). (E) Expression of HIF target genes in HT-29 cells grown in 2% O_2_ ± 100 ppm H_2_S for 24 h normalized to the mean of 3 reference genes (n=4). (F) Scheme showing activation of green fluorescence by the hypoxia sensor UnaG. (G) Effect of H_2_S (0-100 ppm) on hypoxia (2% O_2_) induced fluorescence in UnaG-expressing HT-29 cells. (H) Quantitation of data in G relative to the minus H_2_S condition (n=3). (I) Fluorescence in UnaG-expressing HT-29 cells grown in the presence of 2%-21% O_2_. (J) Dependence of UnaG fluorescence on O_2_ concentrations (normalized to 2% O_2_=1) and fit to an exponential decay curve (n= 3). The green dots correspond to fluorescence observed at the indicated concentration of H_2_S (data from G). Scale bar is 200 μm in all images.

A limitation of bolus Na_2_S treatment is that the concentration of H_2_S is unstable over the course of the experiment due to its volatilization and metabolism. To circumvent this drawback, cells were cultured in a custom growth chamber with an H_2_S atmosphere to mimic chronic exposure experienced by colonocytes (30). The H_2_S concentration was varied between 0-100 ppm for 24 h, which corresponds to a dissolved H_2_S concentration in the culture medium of 0-20 µM and does not affect cell viability (30). A statistically significant decrease in HIF-1α was observed at 25 ppm H_2_S (4 μM dissolved sulfide) and HIF-1α stabilization was completely abolished at 100 ppm H_2_S (20 μM dissolved sulfide) (Fig. 2C,D). RT-qPCR analysis revealed lower mRNA levels of HIF-1α target genes, CA9, GLUT1, HK1 and VEGFA1, consistent with destabilization of the transcription factor at 100 ppm H_2_S (Fig. 2E). Since acute and chronic H_2_S similarly destabilized HIF-1α, we used these regimes interchangeably, depending on whether a relatively fast response (e.g. inhibition of new protein synthesis) or a slower response (e.g., sensor fluorescence) was being monitored.

### H_2_S shifts the threshold for hypoxia sensing

We expressed the genetically encoded fluorescent hypoxia sensors UnaG and dUnaG, which are controlled by HIF-1α-dependent recognition of the hypoxia responsive element (Fig. 2F) (31). UnaG- and dUnaG-expressing HT-29 cells grown at 2% O_2_ exhibited robust fluorescence compared to empty vector controls (Fig. 2G, Supplementary Fig. 3A). In the presence of increasing H_2_S, the decrease in fluorescence paralleled the destabilization of HIF-1α (Fig. 2H, Supplementary Fig. 3D). The sensors were also tested in HEK293 cells, which showed a qualitatively similar response to H_2_S (Supplementary Fig. 3B-C,E-F). The possibility that H_2_S directly quenches fluorescence was ruled out by treating UnaG-expressing HT-29 cells with a large bolus of H_2_S (1 mM), which had no effect on hypoxia-induced fluorescence (Supplementary Fig. 3G,H).

As expected, UnaG and dUnaG fluorescence in HT-29 and HEK293 cells decreased as the O_2_ concentration was increased from 2 to 21% (Fig. 2I,J, Supplementary Fig. 4A-F). The effective intracellular O_2_ sensed in cells grown at 2% O_2_ in the presence of H_2_S could then be estimated from the O_2_ dependency curves of the fluorescent sensors. Even at low concentrations, H_2_S profoundly affected the HIF-1α response and HT-29 cells grown in 2% O_2_ in the presence of 25 or 100 ppm H_2_S reported the equivalent 5 or 15% O_2_, respectively (Fig. 2J, Supplementary Fig. 4D). In contrast, the sensor response at 2% O_2_ ± 10 ppm H_2_S was indistinguishable, indicating that the cellular capacity for sulfide oxidation protected against O_2_ accumulation at this concentration of H_2_S and/or resulted from the limited sensitivity of the sensor to small changes in O_2_ levels. Similar estimates for an H_2_S-dependent increase in effective O_2_ were obtained with HEK293 cells, which reported the equivalent of 9 or ∼14% O_2_ with 25 or 100 ppm H_2_S, respectively (Supplementary Fig. 4E,F, Supplementary Table 2).

### SQOR levels correlate inversely with HIF-1α sensitivity to H_2_S

Comparison of HIF-1α destabilization across four cell lines revealed different sensitivities to H_2_S (Fig. 3A) and raised the question as to whether the differences were correlated with their sulfide oxidation capacity. We found an inverse correlation between SQOR expression levels and HIF-1α destabilization (Fig. 3B,C), consistent with a primary role for this enzyme in regulating intracellular H_2_S and therefore, O_2_ levels (Fig. 1A). Accordingly, the H_2_S effect was exacerbated when SQOR was knocked down (in HT-29^SQOR^ ^KD^), which were slower to regain stable HIF-1α expression compared to scrambled controls (HT-29^Scr^) (Fig. 3D,E).

**Figure 3:**
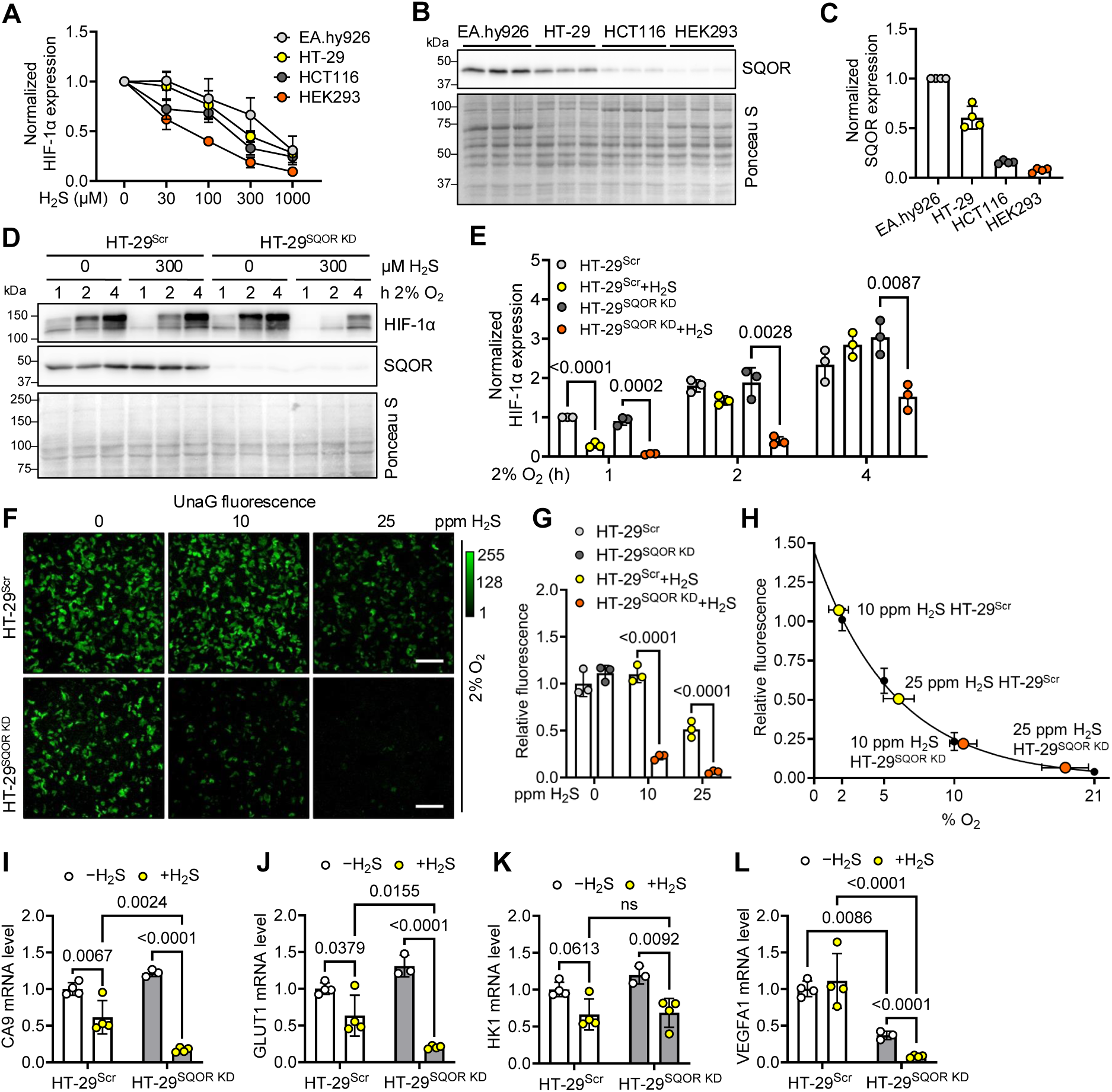
SQOR expression affects sensitivity of HIF-1α to H_2_S. (A) HIF-1α level relative to untreated controls in EA.hy926, HT-29, HCT116, and HEK293 cells grown at 2% O_2_ and varying Na_2_S (0-1 mM). (B) SQOR expression in EA.hy926, HT-29, HCT116, and HEK293 cell lysates detected by western blotting. (C) Quantitation of SQOR levels in B normalized to Ponceau S staining relative to levels in EA.hy926 cells, which have the highest expression (n=4). (D) HIF-1α expression in HT-29^Scr^ versus HT-29^SQOR^ ^KD^ cells in 2% O_2_ treated with 300 µM H_2_S for 1-4 h. (E) Quantitation of HIF-1α levels in D normalized to Ponceau S staining and shown relative to levels in HT-29^Scr^ cells after 1 h (n=3). (F) Effect of H_2_S (10 and 25 ppm, 24 h) on UnaG expression in HT-29^Scr^ and HT-29^SQOR^ ^KD^ cells grown in 2% O_2_. The images are representative of 3 independent experiments. Scale bar is 200 μm. (G) Quantitation of data in F normalized to HT-29^Scr^ without H_2_S. (H) UnaG fluorescence in G was plotted on the standard curve for UnaG fluorescence on O_2_ concentration. (I-L) Expression of HIF target genes in HT-29^Scr^ and HT-29^SQOR^ ^KD^ cells grown in 2% O_2_ ± 25 ppm H_2_S for 24 h. Expression of: (I) CA9, (J) GLUT1, (K) HK1, and (L) VEGFA1 mRNA normalized to the mean of 3 reference genes (n=3 or 4).

UnaG- and dUnaG-expressing HT-29^SQOR^ ^KD^ cells were hypersensitive to low (10 ppm) and moderate (25 ppm) H_2_S exposure resulting in ∼80% and >95% decrease in fluorescence intensity, respectively, compared to control cells (Fig. 3F,G; Supplementary Fig. 5A,B). The fluorescence corresponded to intracellular O_2_ levels of 10-14% and 18-23% for 10 and 25 ppm H_2_S, respectively (Fig. 3H, Supplemental Fig. 5C, Table S2). Further, HT-29^SQOR^ ^KD^ cells exhibited greater HIF-1α destabilization than scrambled controls when grown at moderate H_2_S (25 ppm, 24 h) (Supplemental Fig. 5D,E). These results were mirrored by the suppression of HIF-1α target genes with the exception of HK1 (Fig. 3I-L).

### H_2_S activates PHD to destabilize HIF-1α

The effect of H_2_S on destabilization of HIF-1α could be exerted at or downstream of PHD, which requires 2-ketoglutarate and iron in addition to O_2_ for activity (Fig. 4A). The PHD inhibitor roxadustat or FG4592, induced a dose-dependent (0-100 μM) increase in UnaG fluorescence in cells grown at 21% O_2_, which was insensitive to H_2_S (Supplementary Fig. 6A, Fig. 4B,C). HIF-1α stability was unaffected by acute or chronic H_2_S treatment in the presence of FG4592 (Fig. 4D,E and Supplementary Fig. 6B,C). H_2_S similarly had no effect on HIF-1α stability in the presence of the iron chelator deferoxamine (Fig. 4F-H). These results are consistent with the model that PHD activity is required for H_2_S-dependent HIF-1α destabilization. In the presence of H_2_S and the proteasome inhibitor bortezomib, an increase in the proportion of the ubiquitinated relative to the total HIF-1α pool was seen (Supplementary Fig. 6D-F). A hydroxylated HIF-1α-specific antibody revealed that this pool increased relative to the total HIF-1α pool in bortezomib- and H_2_S-treated cells (Fig. 4I-K), indicating an increase in PHD activity under these conditions.

**Figure 4:**
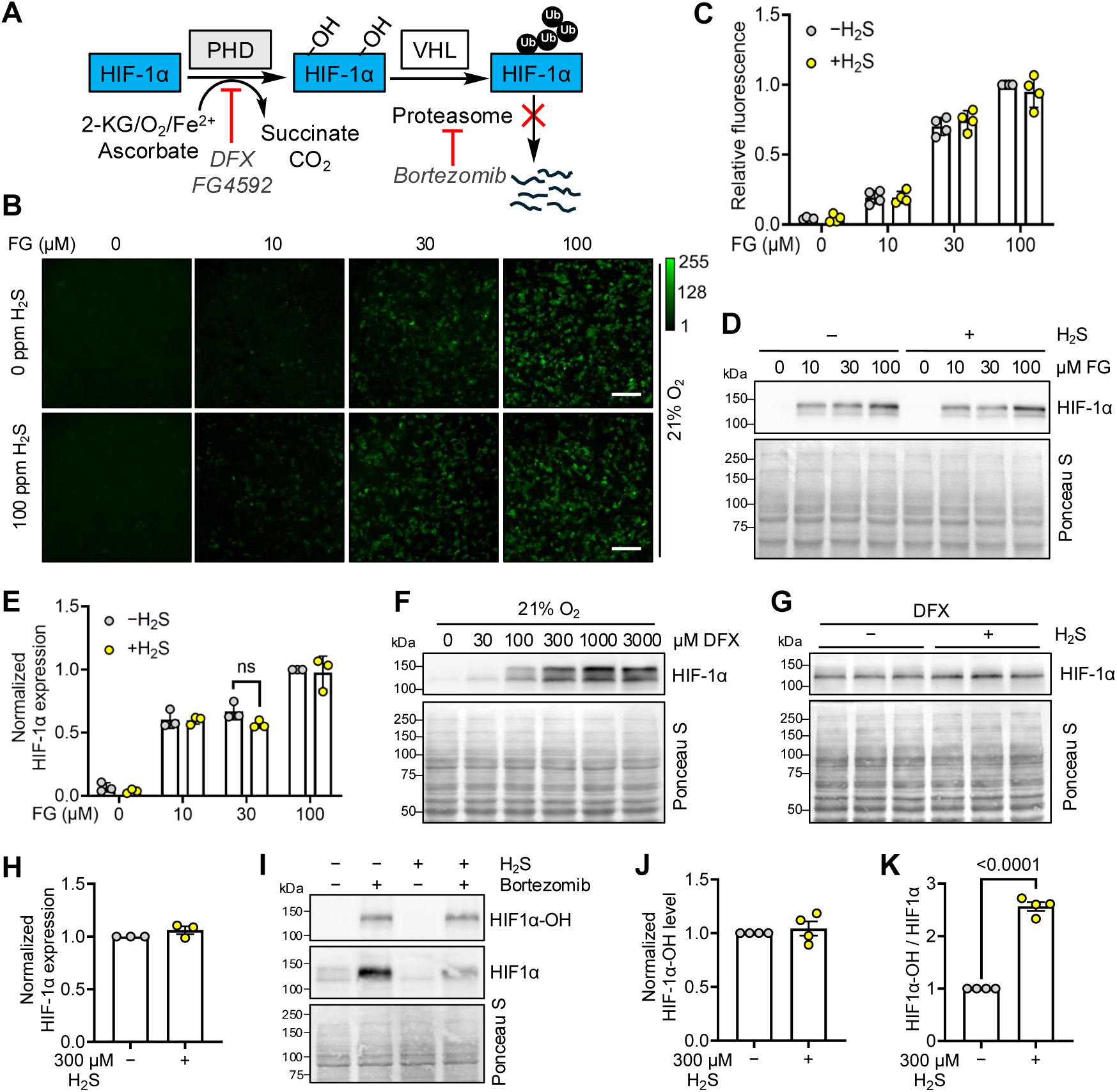
Increased PHD activity mediates hypoxic HIF destabilization by H_2_S. (A) Scheme showing targets of select inhibitors of the O_2_-dependent HIF degradation pathway. KG is α-ketoglutarate. (B) UnaG-expressing HT-29 cells were cultured for 24 h in 21% O_2_ ± 100 ppm H_2_S in the presence of 0, 10, 30, or 100 μM FG4592 (FG). Scale bar is 200 μm. (C) Quantitation of data in B expressed relative to the 100 μM FG4592 minus H_2_S condition (n=4). (D) HIF-1α expression in HT-29 cells treated with 0, 10, 30, and 100 μM FG4592 and ± 100 ppm H_2_S (E) Quantitation of data in D normalized to Ponceau S staining (n=3). (F) Western blot analysis of HIF-1α in HT-29 cells grown in 21% O_2_ and treated with varying concentrations of the PHD inhibitor deferoxamine (DFX). (G) HIF-1α expression in HT-29 cells grown in 21% O_2_ ± 300 µM DFX and 300 µM Na_2_S for 1 h. (H) Quantitation of data in G normalized to Ponceau S staining (n=3). (I) HT-29 cells were pre-treated with bortezomib for 6 h and then grown in 2% O_2_ ± 300 µM Na_2_S. The hydroxylated form of HIF-1α was detected by western blot analysis. (J,K) Quantitation of data in I normalized to Ponceau S staining (J) or total HIF-1α (K) (n=4).

In principle, H_2_S could additionally destabilize HIF-1α levels by activating lysosomal degradation (32) or inhibiting new protein synthesis (33) (Supplementary Fig. 7A). Both mechanisms were, however, ruled out. The lysosomal inhibitor chloroquine increased HIF-1α levels in cells grown at 2% O_2_ but was unaffected by H_2_S, and puromycin, which allows monitoring of nascent polypeptide synthesis, was unaffected by acute or chronic H_2_S (Supplementary Fig. 7B-F). Collectively, these results support a model in which H_2_S stimulates substrate (i.e., O_2_) level activation of PHD to enhance HIF degradation. Further, eIF2α phosphorylation was unaffected by sulfide (Supplementary Fig. 7G-J), as reported previously (20), which is in contrast to H_2_S-dependent enhancement of phospho-eIF2α in mouse embryonic fibroblast and HeLa cells (34). It is unclear whether these differences reflect differences in cell lines and/or or stem from other mechanistic differences such as H_2_S-dependent inhibition of protein phosphate PP1c, in some but not other cells.

### *Lb*NOX counters the effect of H_2_S on HIF-1α

As a further test of our hypothesis that H_2_S destabilizes HIF-1α by increasing intracellular O_2_ levels, we used *Lactobacillus brevis* NADH oxidase (*Lb*NOX), to promote O_2_ consumption (35) by dissipating the NADH pool (Fig. 5A). Mitochondrial but not cytoplasmic expression of *Lb*NOX enhanced HIF-1α stabilization and attenuated H_2_S-induced HIF-1α destabilization when the sulfide concentration was low (25 ppm, 24 h in 2% O_2_) (Fig. 5B,D). However, when the sulfide concentration was raised to 100 ppm, the effect of mitochondrial *Lb*NOX on HIF-1α stability was significantly diminished (Fig. 5C,E).

**Figure 5:**
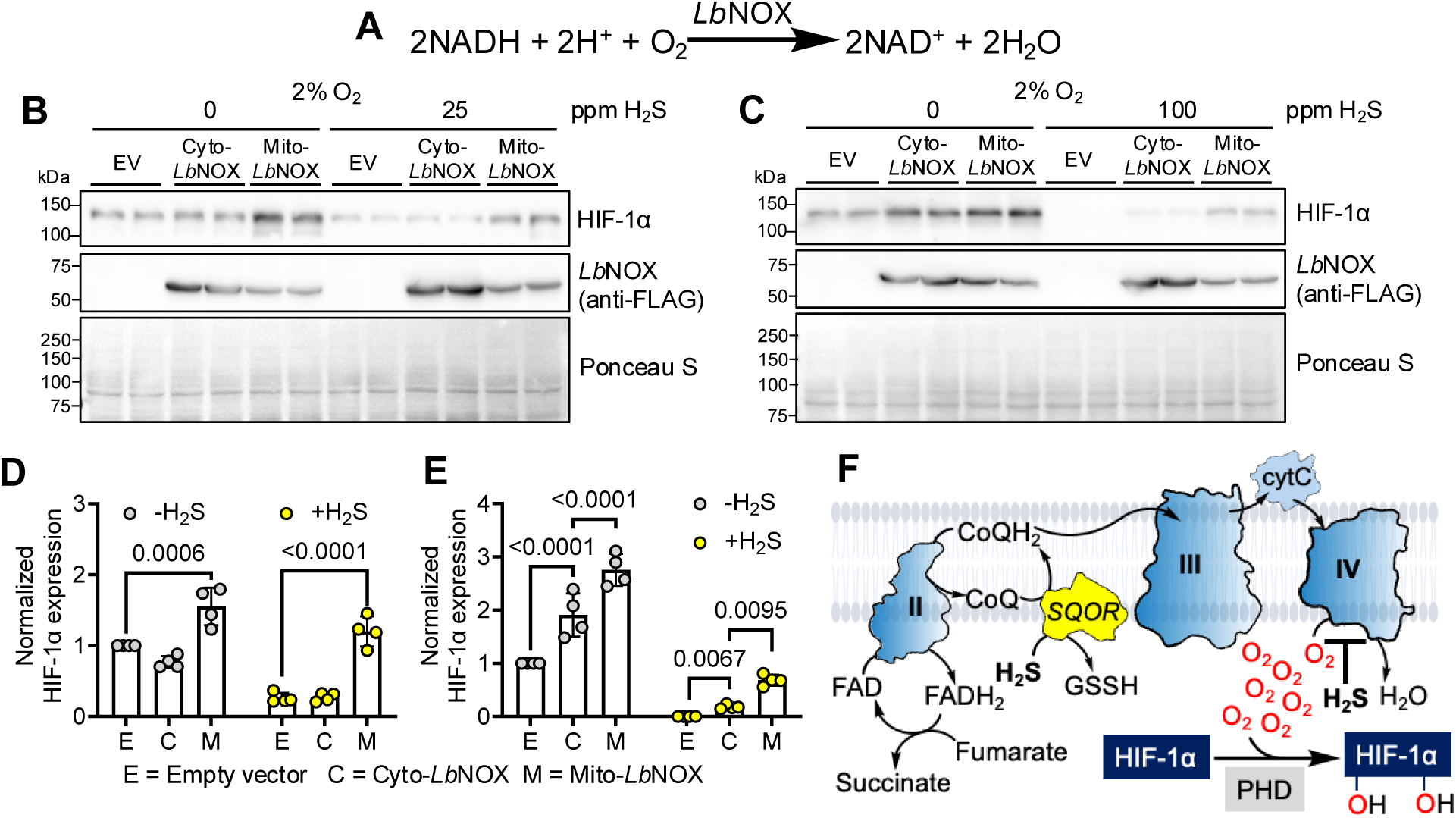
*Lb*NOX restores HIF under H_2_S. (A) O_2_ consumption reaction catalyzed by *Lb*NOX. (B,C) Western blot analysis of HIF1α and *Lb*NOX expression in HT-29 cells transfected with empty vector (EV), cytosolic (Cyto-*Lb*NOX) or mitochondrial (Mito-*Lb*NOX) *Lb*NOX, and cultured in 2% O_2_ ± 25 ppm H_2_S (B) or ± 100 ppm H_2_S (C) for 24 h. (D,E) Quantitation of HIF-1α data in B (D) and C (E) normalized to Ponceau S staining (n=4). (F). O_2_ and fumarate serve as terminal electron acceptors for H_2_S oxidation, allowing sulfide concentration to dynamically regulate intracellular O_2_ levels.

## DISCUSSION

The cartography of intracellular O_2_ is dynamically regulated by mitochondrial respiration and is important for protecting against O_2_ cytotoxicity (36, 37). Consequently, mitochondrial diseases are characterized by dysregulated O_2_ metabolism and venous hyperoxia during exercise (38) and interestingly, show promise for hypoxia therapy (39). In this study, we establish H_2_S as a powerful regulator of intracellular O_2_, which has important implications for O_2_ sensing and redox metabolism especially in tissues with low physiological O_2_.

Using a custom sulfide growth chamber to simulate chronic H_2_S exposure (40), we demonstrated that even a low concentration of H_2_S (25 ppm in the atmosphere or 4 µM in culture medium) induces a profound shift such that 2% O_2_ elicits a response equivalent to 5-9% ambient O_2_ (Fig. 2J and Supplementary Table 2). H_2_S exhibits a dose-dependent modulation of O_2_ tension that is sensitive to SQOR levels across cell lines (Fig. 3A-C) and is consistent with its contribution to a high intracellular H_2_S turnover rate (41, 42). We postulate that SQOR activity in response to H_2_S biogenesis or exogenous exposure from gut lumen regulates ETC flux and therefore, O_2_ metabolism in healthy tissue (43, 44). The physiological significance of these observations is underscored by the importance of SQOR for endothelial cell proliferation under hypoxic conditions, post-ischemia neovascularization, tumor neoangiogenesis and rescue from hypoxic brain injury (45, 46).

SQOR is ubiquitously expressed, and based on the human protein atlas database, its highest levels are found in the gastrointestinal tract, skeletal muscle, nasopharynx and kidney. Whole-body SQOR-deficient mice cease to grow at weaning and die within 8-10 weeks of age (46, 47), indicating severe growth impairment in the absence of H_2_S oxidation capacity. Inherited SQOR deficiency presents as Leigh syndrome (48), a disease with over 75 contributing genetic loci, which is characterized by profound neurological and neuromuscular defects (49). In contrast to murine models, the clinical presentation in the limited number of SQOR deficiency patients described so far, has ranged from 4-8 years of age, possibly suggesting incomplete penetrance of the associated variants (48).

H_2_S-dependent regulation of O_2_ consumption and accumulation is paradoxical since the latter comes at the cost of inhibiting respiration and decreasing O_2_-dependent sulfide clearance. However, sulfide consumption and O_2_ accumulation can occur simultaneously due to the flexibility inherent in the ETC for coupling H_2_S oxidation to fumarate versus O_2_ reduction (50, 51) (Fig. 5F). We posit that physiologically relevant hypoxia modulation can occur within an H_2_S concentration window where complex IV is partially inhibited, and contribute to regional O_2_ contours that can vary significantly even within a given tissue (17, 52). The durability of the complex IV response to fractional inhibition even at low H_2_S (53), further suggests that the effect of sulfide on O_2_ accumulation might be long-lived. The physiological importance of sulfide-dependent O_2_ regulation is revealed by the loss of Fe-S proteins in colon of mice lacking SQOR in intestinal epithelial cells (Fig. 1H,I). We presume that damage accrues at O_2_ and ROS-sensitive sites as H_2_S levels, and therefore O_2_ levels, rise. We speculate that on the other hand, the beneficial boost in exercise endurance by the natural product ergothioneine, which increases endogenous H_2_S biogenesis (54, 55), could be due in part to increased O_2_ availability across skeletal muscle.

Mitochondria function as O_2_ sinks (56) and help establish intracellular O_2_ gradients (57–59). Perinuclear mitochondria decrease nuclear O_2_ exposure and limit its genotoxicity (60). The effectiveness of mitochondrial but not cytoplasmic *Lb*NOX to rescue hypoxia sensing was dependent on H_2_S concentration, further illuminating the interplay between O_2_ and H_2_S and its sensitivity to their respective concentrations. Decreased ETC flux due to H_2_S leads to a reductive shift in the NAD pool(25), which can be dissipated by *Lb*NOX that concomitantly utilizes O_2_ (Fig. 5A). Since HIF is a cytoplasmic protein, it is curious that only mitochondrially-targeted *Lb*NOX stabilized HIF-1α, which could be explained in part by its higher O_2_ consumption (35). Additionally, by increasing coenzyme Q availability, mito-*Lb*NOX activity could enhance SQOR-dependent H_2_S clearance, decreasing complex IV inhibition. Importantly, in addition to PHD (*K_M_*_(O2)_ =67-85 µM (61)), other enzymes that exhibit varying degrees of unsaturation at ambient intracellular O_2_ (62) are also expected to be sensitive to H_2_S-mediated changes in O_2_. Some notable examples include the KDM family of lysine demethylases that use histone and non-histone substrates and the TET enzymes that catalyze hydroxylation of 5-methyl cytosine on DNA. These functions are central to epigenetic regulation and other processes such as mitochondrial biogenesis(62).

In summary, we demonstrate that H_2_S shifts the threshold for hypoxia sensing across cell lines by increasing intracellular O_2_. Chronic H_2_S exposure under ambient hypoxia induces an oxidative shift in the mitochondrial cysteine proteome and destabilizes Fe-S proteins involved in oxidative energy metabolism in cells. In a physiological setting, it is the loss of SQOR in gut epithelium that renders the tissue vulnerable to oxidative changes and reveals that sulfide oxidation capacity buffers O_2_ cytotoxicity. Perturbed O_2_ metabolism is likely to underlie the pathologies associated with H_2_S overload such as SQOR (48), ETHE1 (63) or sulfite oxidase (64) deficiency and our study provides a foundation for investigating these links in these and other diseases including acute respiratory distress syndrome and inflammation that are characterized by hyperoxia. Finally, increasing H_2_S synthesis or decreasing its oxidation, might be beneficial in diseases characterized by hypoxia, including metabolic and cardiovascular diseases.

*Limitations of our study*. While our study provides a quantitative evaluation of the magnitude of H_2_S-induced perturbation of the intracellular O_2_ economy in the setting of 2-10% ambient O_2_, the limited dynamic range of the UnaG and dUnaG hypoxia sensors restricted their utility outside this O_2_ concentration range. Second, while the *in vivo* mouse colon data are consistent with H_2_S-dependent dysregulation of O_2_ metabolism, our experiments were largely focused on cell lines adapted to long-term culture under normoxic conditions and additionally, did not examine HIF-2α modulation. Finally, our study assessed the effect of exogenous H_2_S at low O_2_ to simulate the environment in the colon lumen. Endogenous upregulation of H_2_S biogenesis via diet (e.g., increased methionine and/or cysteine) and/or via overexpression of the H_2_S-synthesizing enzymes would broaden the scope of this study and uncover its regulation in other contexts.

## METHODS

### Materials

RPMI 1640 medium (11875093), RPMI+HEPES (22400089), DMEM (11995065), fetal bovine serum (A5256701), 1x pen-Strep (15140122), geneticin (10131035), trypsin-EDTA (25300062), and phosphate buffered saline (10010023) were from Gibco. Sodium sulfide nonahydrate (431648), dimethyl sulfoxide (D2650), chloroquine (C6628), protease inhibitor (P8340), doxycycline (D3447) and puromycin (P8833) were from Sigma-Millipore. Methanol (A4544) and Tween-20 (BP337500) were from Fisher. PVDF membranes (1620177), blot filter paper (1703932) and Clarity Western ECL Substrate (102032117 and 102032118) were from Bio-Rad. Deferoxamine (14595), bortezomib (10008822), and FG4592 (15294) were from Cayman. Phosphatase inhibitor cocktail (HY-K0021) was from MedChemExpress. Nonidet P40 (74385) was from Fluka BioChemika. Gas cylinders of H_2_S (500 L 5000 ppm H_2_S in N_2_ within 1% accuracy), normal air containing CO_2_ (21% O_2_, 5% CO_2_, and 74% N_2_) and hypoxic air (2% O_2_, 5% CO_2_, 93% N_2_) were from Cryogenic Gases (Detroit MI, USA).

The primary antibodies used in this study were: anti-HIF1α (Abcam ab179483, 1:3000), anti-HIF1α-OH Pro-564 (CST 3434, 1:3,000), anti-MT-CO2 (Abcam ab110258, 1:5,000), anti-SQOR (Proteintech 17256-1-AP, 1:5,000), anti-puromycin (DSHB PMY-4A4, 1:500), anti-eIF2a (CST 9722, 1:3,000), anti-p-eIF2a phospho-S51 (Abcam ab32157, 1:3,000), anti-NDUFS8 (Abcam ab170936, 1:1,000), anti-ACO2 (Abcam ab129069, 1:10,000), anti-LIAS (Abcam ab246917, 1:1,000), anti-UQCRSF1 (Abcam ab14746, 1:1,000), anti-FLAG (Sigma F1804, 1:5,000) and anti-lipoic acid (Abcam ab58724, 1:1,000). Secondary antibodies conjugated to HRP were anti-rabbit (Abcam ab6721) and anti-mouse (Abcam ab205719) used at a 1:10,000 dilution.

### Cell culture

HT-29, HCT116, HEK293, and EA.hy296 cells were obtained from ATCC. HT-29^Scr^, HT-29^SQORKD^, HT-29 cyto-LbNOX, HT-29 mito-LbNOX, and HT-29 EV cells were previously generated in the laboratory (25, 29). HT-29 HRE-UnaG and HT-29 HRE-dUnaG stable cells were generated in this study as described below. HT-29 cells were cultured in RPMI 1640 medium supplemented with 10% FBS and 1x Pen-Strep. All other cell lines were cultured in DMEM with 10% FBS and 1x Pen-Strep. HT-29 cells stably expressing cyto-*Lb*NOX, mito*Lb*NOX, empty vector, HRE-UnaG, and HRE-dUnaG were supplemented with 300 μg/mL geneticin. HT-29^SQORKD^ and HT-29^Scr^ were supplemented with 1 μg/mL puromycin. All cell lines were cultured in a humidified cell culture incubator at 37 °C with 5% CO_2_ and ambient air except in hypoxia experiments, which were conducted at 2% O_2_, 93% N_2_, and 5% CO_2_ unless otherwise noted. During 24 h hypoxia experiments, cells were switched to fresh medium containing HEPES at twice the normal volume to minimize acidification (30).

### Generation of stable cell lines for HRE-UnaG and HRE-dUnaG

Plasmids for expression of the UnaG and dUnaG fluorescent proteins under control of a hypoxia responsive element (HRE) promoter were obtained from Addgene (plasmid #s 201710 and 201711). Transfection-grade plasmids were purified from *E. coli* (Qiagen 12123) and the insert region was confirmed by sequencing (Eurofins). HT-29, HT-29^Scr^, and HT-29^SQOR^ ^KD^ cells were transfected with the HRE-UnaG and HRE-dUnaG expression plasmids in 24-well culture plates. After 24 h, cells were switched to fresh medium containing geneticin at 300 μg/mL. Cells were incubated in the presence of geneticin for 2-3 weeks, after which all non-transfected cells had died. Surviving cells were expanded in 6-well plates and a portion of the cells were split evenly into two 24-well plates for a trial experiment. One plate was left in normal cell culture conditions and the other incubated under hypoxia for 24 h. The fluorescence in each well was quantitated using an EVOS microscope (EVOS M5000) and the colonies with the highest fluorescence for each plasmid were expanded to make cell stocks.

### Western blotting

#### Sample preparation

Cells were seeded 24 h prior to the start of experiments in 35 mm or 60 mm plates such that the confluency at collection was 70-80%. Fresh medium was added at the start of experiments. For detection of protein expression, cell plates were washed with ice-cold PBS and then scraped with a cell scraper on ice in lysis buffer (20 mM HEPES pH 7.4, 25 mM KCl, 0.5% Nonidet P40, 1x protease inhibitor) and immediately frozen on dry ice. Samples were stored at -80 °C until further use.

#### Development of SDS-PAGE western blots

Frozen lysates were rapidly thawed in a 37 °C water bath for 50 sec until the ice was nearly melted and then vortexed for 15 sec and immediately centrifuged at 13,000 x *g* for 5 min at 4 °C. Cleared lysates were transferred to new tubes and immediately denatured with 4x SDS+DTT buffer (200 mM Tris-HCl pH 6.8, 40% glycerol, 8% SDS, 0.1% bromophenol blue, 400 mM DTT) and incubated for 5 min at 80 °C. In parallel, 4 μL cleared lysate per sample was used for protein assay with 1 mL Bradford reagent (Bio-Rad). Samples (30 μg per lane) were run on 8, 10, or 12% Tris-glycine SDS gels at 100-130 V, transferred to PVDF membranes, and blocked for 30 min with 5% milk in Tris buffered saline containing 0.1% Tween 20 (TBST). Membranes were probed overnight with primary antibodies diluted in 5% milk and then washed 3 x 10 min with TBST. Horseradish peroxidase (HRP)-linked secondary antibodies were incubated for 1-2 h with the membranes followed by another round of 3 x 10 min washes. Blots were developed with ECL substrate and imaged with a Bio-Rad ChemiDoc imaging system. Equal loading was verified by Ponceau S staining of membranes. Blot image files were quantitated by pixel intensity in ImageJ and normalized to the quantified Ponceau S staining. When two bands were seen with the HIF-1α antibody, the intensities of both were combined for quantitation.

### Intracellular O_2_ assay

HEK293 cells were plated at 50% confluency in 6-well plates. After 24 h to allow attachment to the plate, cells were transfected with HRE-UnaG, HRE-dUnaG or empty vector with lipofectamine 3000 (Invitrogen L3000001). After 24 h of transfection, cells were split evenly between three 24-well plates. After another 24 h to allow re-attachment of cells, 1 mL fresh medium was carefully added to each well and the plates were incubated in the following atmospheres: (i) A baseline hypoxia control (2% O_2_, 5% CO_2_, 93% N_2_); (ii) Hypoxia as in (i) admixed with 10, 25, or 100 ppm H_2_S as described previously (30); and (iii) A calibration control at either 5%, 10%, or 21% O_2_, 5% CO_2_, with the balance being N_2_. After 24 h of incubation, all plates were imaged using the EVOS microscope at 20x magnification using GFP filter and brightfield to obtain representative images. Images were quantitated using ImageJ. A calibration bar denoting the pixel intensity range is provided in each figure. For HT-29 cells stably expressing HRE-UnaG and HRE-dUnaG, a similar procedure was followed except that the transient transfection steps were omitted. Instead, cells were plated directly in 3 x 24-well plates at ∼40% confluency 24 h prior to switching to hypoxia.

### HIF stability assays

#### Bolus H_2_S treatment

HT-29, HEK293, HCT116, and EA.hy296 cells were plated in 6 cm dishes and grown to 70-80% confluency. On the day of the experiment, fresh medium supplemented with a bolus of Na_2_S was added and cells were immediately transferred to the hypoxia chamber. HIF was detected from the cell lysates and normalized to Ponceau S staining.

#### H_2_S time course and dose response assays

For the time course experiments, HT-29 cells (70-80% confluency) were cultured in 6 cm plates with fresh growth medium and treated with either vehicle or 300 μM Na_2_S and then immediately placed in the hypoxia chamber. Plates were removed from the chamber for lysate collection at intervals ranging from 0.25 to 4 h. For dose response experiments, cells treated with varying doses of Na_2_S (0-1000 μM) were incubated under 2% hypoxia for 1 h. HIF was detected in the cell lysates and blots were normalized by Ponceau S staining. Na_2_S-treated cells were cultured separately from non-Na_2_S treated cells to prevent contamination of aerosolized H_2_S in the culture medium of the control samples.

#### PHD inhibitor treatments

HT-29 cells were cultured under normoxic conditions for 1 h in medium treated with 0, 30, 100, 300, 1000, or 3000 μM deferoxamine (DFX) or with 0, 3, 10, 30, 100, or 300 μM FG4592 dissolved in 0.1% DMSO. HIF was detected in cell lysates and normalized to Ponceau S staining.

#### HIF ubiquitination and proline hydroxylation assays

HT-29 cells were pre-treated with the proteasome inhibitor bortezomib for 6 h and then with a bolus of Na_2_S and incubated under hypoxia for 1 h. Cell lysates were isolated, run on 8% gels and probed with anti-HIF-1α to observe high molecular weight bands corresponding to ubiquitinated HIF. Blots were probed with Anti-HIF-1α-OH to detect the proline hydroxylated form. Ubiquitinated or hydroxylated HIF-1α were normalized to total HIF-1α levels and to Ponceau S staining.

#### Chloroquine assay for lysosomal degradation

HT-29 cells were treated with 50 μM chloroquine to inhibit lysosomal degradation in combination with Na_2_S and incubated for 1 h under hypoxia. HIF stability was assessed in cell lysates and normalized to Ponceau S staining.

#### LbNOX assays

HT-29 cells expressing *Lb*NOX and mito-*Lb*NOX (29) were plated in 6-cm plates and *Lb*NOX expression was induced with 300 ng/mL doxycycline for 24 h. Cells were moved to the hypoxia chamber and exposed to either 0, 25 or 100 ppm H_2_S for 24 h then harvested for protein immunoblotting.

### Puromycin incorporation assay

Cells were switched to medium containing 1 μM puromycin and co-incubated for 1 h with 300 μM Na_2_S. Cell lysates were probed with anti-puromycin antibody to detect puromycin-conjugated nascent peptides as a marker of new protein synthesis. A negative control was run without puromycin. Western blots were normalized to Ponceau S staining.

### RT-qPCR analysis

RNA was extracted from 70-80% confluent cells using the Trizol reagent following the manufacturer’s protocol, and reverse transcribed with murine leukemia virus using random primers. Samples were analyzed by qPCR using SYBR Select Master Mix and primers listed in Supplementary Table 3. All data were normalized to the mean of 3 reference genes, *GUSB*, *TBP*, and *POLR2A*.

### Lipid peroxide assay

All FACS analyses were conducted using the Bio-Rad Ze5 multi-laser, high speed cell analyzer operated with the Everest software package at the University of Michigan Flow Cytometry Core Facility. All data were analyzed using FlowJO (v10.8.1).

#### Lipid ROS analyses using the BODIPY lipid peroxide sensor

HT29 cells were split at 2 million cells per 6 cm plate, cultured overnight, and then had their medium changed to 8 mL before plating for 24 h culture ± 100 ppm H_2_S. Two hours before harvest, positive control cells were treated with 100 µM cumyl hydroperoxide, which was prepared fresh in absolute ethanol as a 1:100 dilution of a 100 mM, 80% technical grade stock (Fisher, AAL0686622). A vehicle control of 10 µL absolute ethanol was included. After 24 h ± 100 ppm H_2_S, the cells were washed twice with 1 mL ice-cold PBS, trypsin digested, pelleted at 1000 x *g* for 5 minutes at 4 ⁰C, and then resuspended in 3 mL of ice-cold blocking buffer (10% FBS in PBS). The cells were transferred to 2 x 1.5 mL tubes. Each tube was centrifuged at 1000 x *g* for 5 min, and then resuspended in prewarmed blocking buffer containing DMSO vehicle or 2 µM of BODIPY and allowed to stain in the dark for 30 min at 37 ⁰C. The DMSO unstained samples were not included in the results but were used to confirm high signal to noise. After staining, the cells were centrifuged at 1000 x *g* for 5 min, resuspended in 2% FBS in PBS, then filtered through 5-mL BD round bottom falcon tubes with cell-strainer caps for FACS analysis on the Bio-Rad Ze5 multi-laser. FlowJO (v10.8.1) was used to conduct gating and ratio the PE and FITC fluorescence emission spectra for each analyzed cell, to plot the histograms of the results, and to calculate the median fluorescence ratios.

### PL-OxICAT analysis of the cysteine redox proteome

HEK293T cells expressing PDK1-TurboID were split at 10% confluency into 10 cm plates and cultured for 2 days before changing medium and growing ± 100 ppm H_2_S for an additional 24 h. Then, each plate was treated with 500 µM biotin in DMSO for 1 h before washing 3x with cold DPBS. Cells were harvested by scraping in DPBS, pelleting at 1600 x *g* for 5 min, aspirating the supernatant, and frozen in liquid nitrogen. Cells were 90-95% confluent at the time of harvest. Frozen cells were then subjected to the redox proteomics workflow as described (27).

### Statistics

All data are presented as mean ± SD with the individual data points displayed. For western blot and fluorescence quantitation, control group data were set to 1 and other data points were displayed as relative levels compared to the controls. Statistical analyses were performed with GraphPad Prism 10.2.2. Unpaired, two-tailed t-tests were used for binary comparisons. One-way ANOVA with Bonferroni’s correction was used for multiple comparisons. The exact p values were displayed above each indicated comparison. Not significant (ns) indicates a p value >0.05.

## ACKNOWLEDGEMENTS

This work was supported in part by the grants from the National Institutes of Health (GM130183 to RB and R35GM134964 to EW).

## DATA AND MATERIALS AVAILABILITY

All data generated and analyzed in this study are included in the main text and Supplementary Information file.

## AUTHOR CONTRIBUTIONS

JB, APL, RS, YS and RB conceptualized the study, JB performed and analyzed the majority of the experiments and was assisted by: DAH for lipid peroxidation assays, DAH and RK for Fe-S protein analysis in cells (DAH) and tissue (RK); DAH, QP and EW for cysteine redox proteomics analysis; APL and RS for HIF analysis. JB and RB drafted the manuscript, and all authors edited and approved the final version.

## Supplementary Table of Content

**Supplementary Table 1.** Mass spectrometric data from PL-OxICAT analysis

**Supplementary Table 2**. Perturbation of intracellular O_2_ levels by H_2_S

**Supplementary Table 3.** List of primers used for RT-qPCR analysis

**Supplementary Figure 1.** H_2_S increases lipid oxidation

**Supplementary Figure 2.** HIF destabilization by H_2_S across cell lines

**Supplementary Figure 3.** H_2_S-induced increase in O_2_ monitored by dUnaG

**Supplementary Figure 4.** O_2_ calibration curves in HEK cells

**Supplementary Figure 5:** SQOR deficiency sensitizes cells to HIF destabilization by H_2_S

**Supplementary Figure 6.** HIF destabilization by H_2_S is PHD-dependent

**Supplementary Figure 7.** H_2_S does not affect HIF-1α synthesis or its lysosomal degradation

**Supplementary Table 2.** Perturbation of intracellular O_2_ levels by H_2_S in HT-29 and HEK cells grown in 2% O_2_ and exposed to the indicated concentrations of sulfide.

| Cell Line | [H<sub>2</sub>S], ppm | % O<sub>2</sub>, Intracellular<sup>1</sup> |  |
| --- | --- | --- | --- |
|  |  | UnaG | dUnaG |
| HT-29 | 10 | 2.0 ± 0.4 | 1.9 ± 0.2 |
| HT-29 | 25 | 5.2 ± 0.2 | 6.3 ± 1.1 |
| HT-29 | 100 | 15.1 ± 3.5 | 17.2 ± 1.1 |
| HEK293 | 10 | 2.0 ± 0.1 | 2.0 ± 0.2 |
| HEK293 | 25 | 8.5 ± 1.4 | 9.5 ± 1.3 |
| HEK293 | 100 | 14.2 ± 4.8 | 13.5 ± 5.2 |
| HT-29<sup>Scr</sup> | 10 | 1.6 ± 0.5 | 1.9 ± 0.7 |
| HT-29<sup>Scr</sup> | 25 | 5.9 ± 1.0 | 5.2 ± 0.8 |
| HT-29<sup>SQOR KD</sup> | 10 | 10.8 ± 0.8 | 14.0 ± 0.6 |
| HT-29<sup>SQOR KD</sup> | 25 | 18.3 ± 1.6 | 23.6 ± 1.7 |

<sup>1</sup>O<sub>2</sub> levels were estimated relative to 2% ambient O<sub>2</sub> and 0 ppm H<sub>2</sub>S

**Supplementary Table 3.**
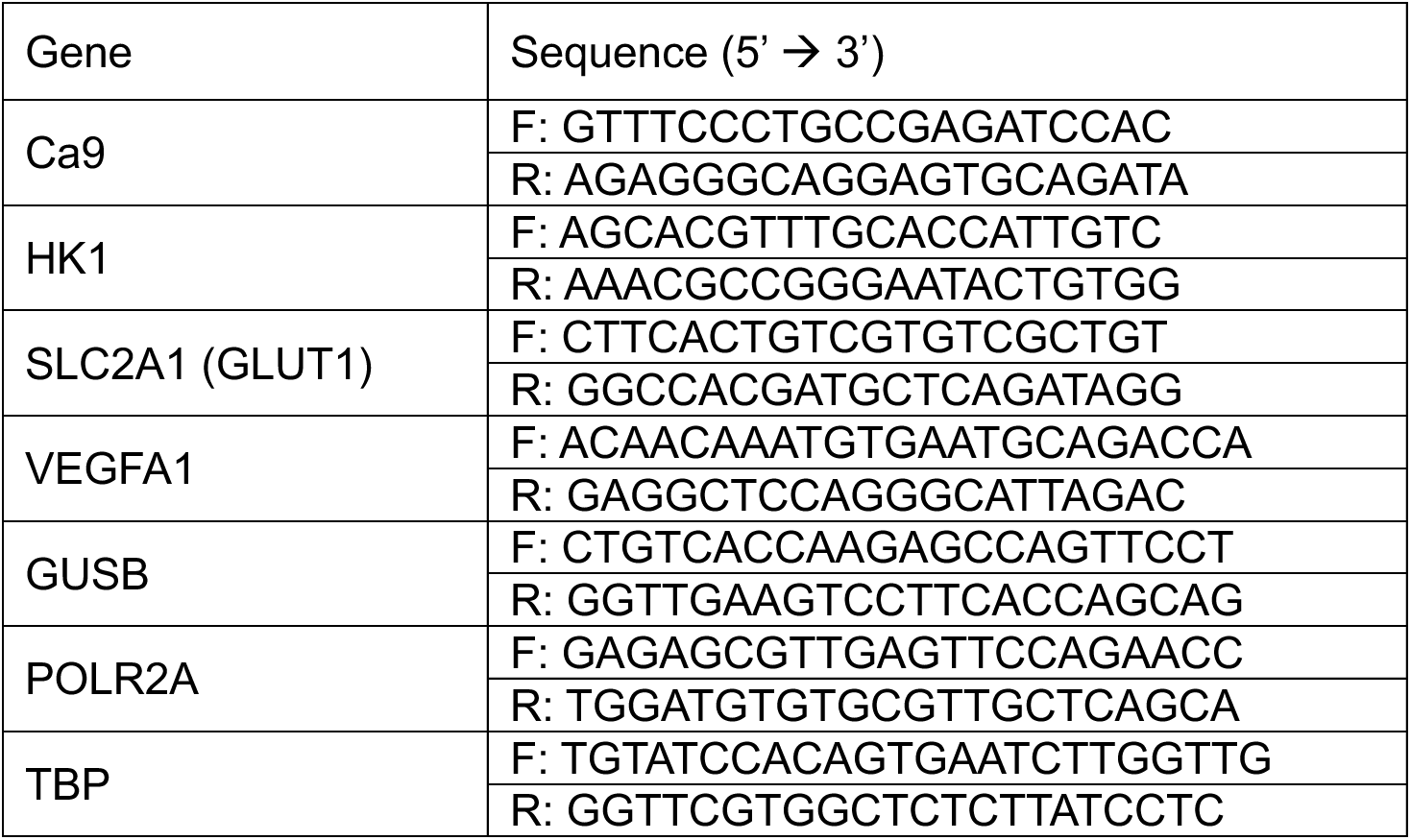
List of primers used for RT-qPCR analysis.

**Supplementary Figure 1:**
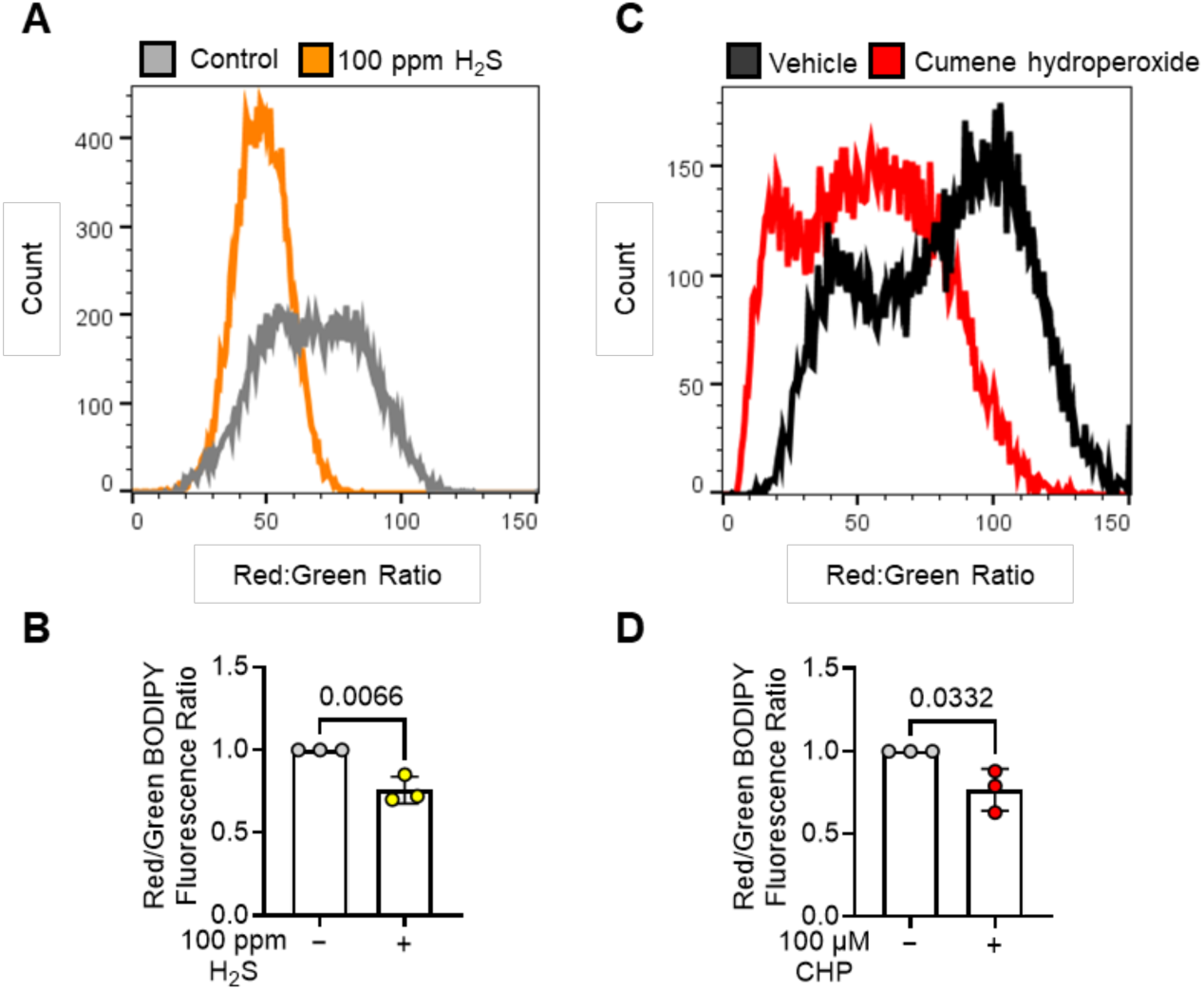
H_2_S increases lipid oxidation. (A) Ratio of red:green fluorescence, indicating the reduced:oxidized ratio of BODIPY 581/591 C11 lipid peroxidation sensor in HT-29 cells exposed to 100 ppm H_2_S for 24 h. (B) Quantitation of the red/green BODIPY fluorescence ratio from A. (C) Positive control with the same cells exposed to 100 µM cumene hydroperoxide (CHP) to induce lipid ROS. (D) Quantitation of the red/green BODIPY fluorescence ratio from C.

**Supplementary Figure 2:**
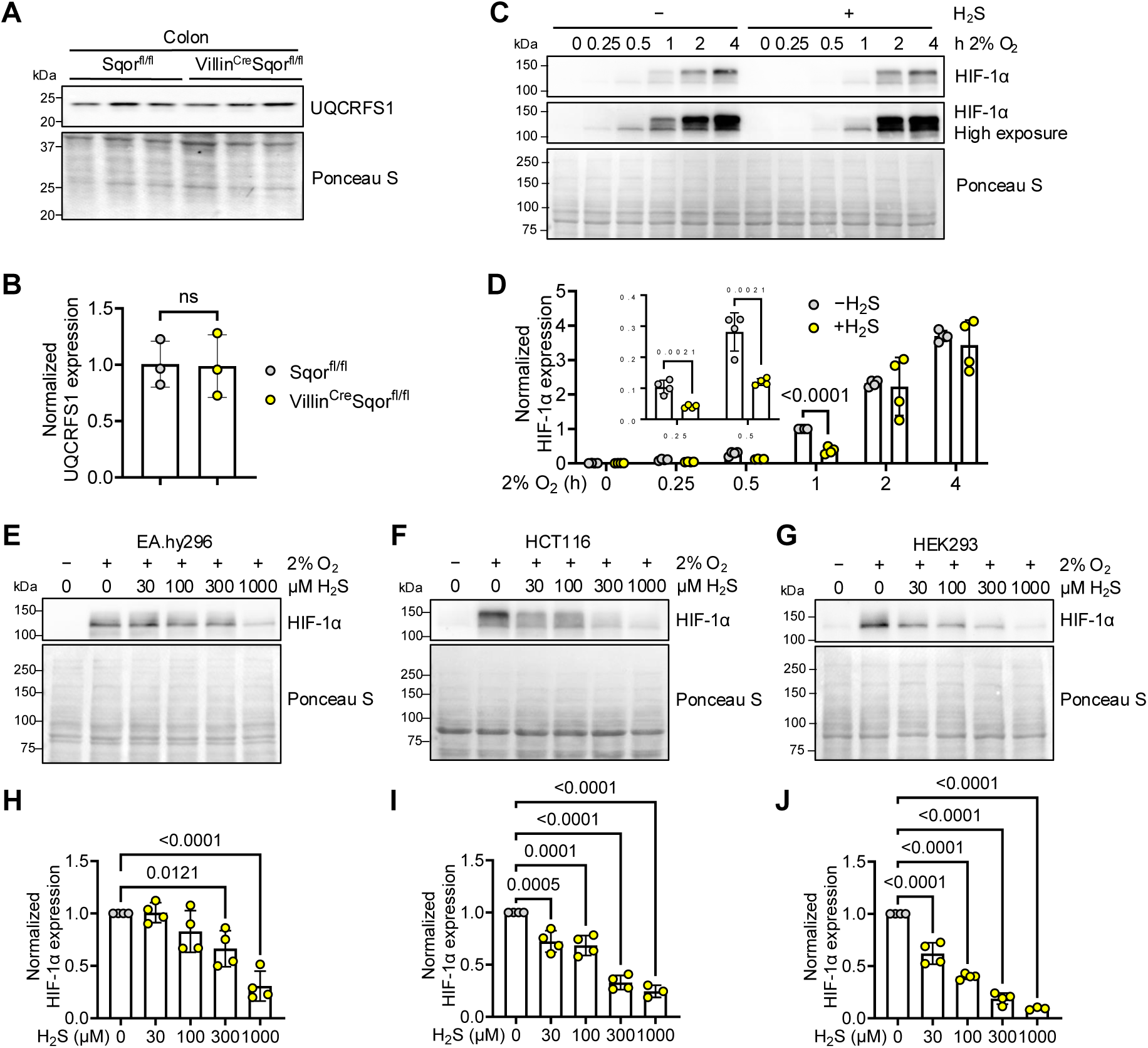
HIF destabilization by H_2_S across cell lines. (A,B) Representative western blots (A) and quantitation (B) of colon UQCRFS1 from Sqor^fl/fl^ (n=3) and *Villin*^Cre^ Sqor^fl/fl^ mice (n=3). (C) HT-29 cells grown in a 2% O_2_ atmosphere were exposed to 300 µM Na_2_S for the indicated times and HIF1α was detected in cell lysates by western blot analysis. (D) Quantitation of HIF-1α intensity in (C) normalized to Ponceau staining and expressed relative to the sample intensity at 1 h without H_2_S. (E-G) H_2_S dose dependence of HIF-1α destabilization in (E) EA.hy296, (F) HCT116, and (G) HEK293 cells grown in 2% O_2_. (H-J) Quantitative analysis of western blots for HIF-1α in E-G, respectively normalized to Ponceau staining and presented relative to untreated controls (n=4).

**Supplementary Figure 3.**
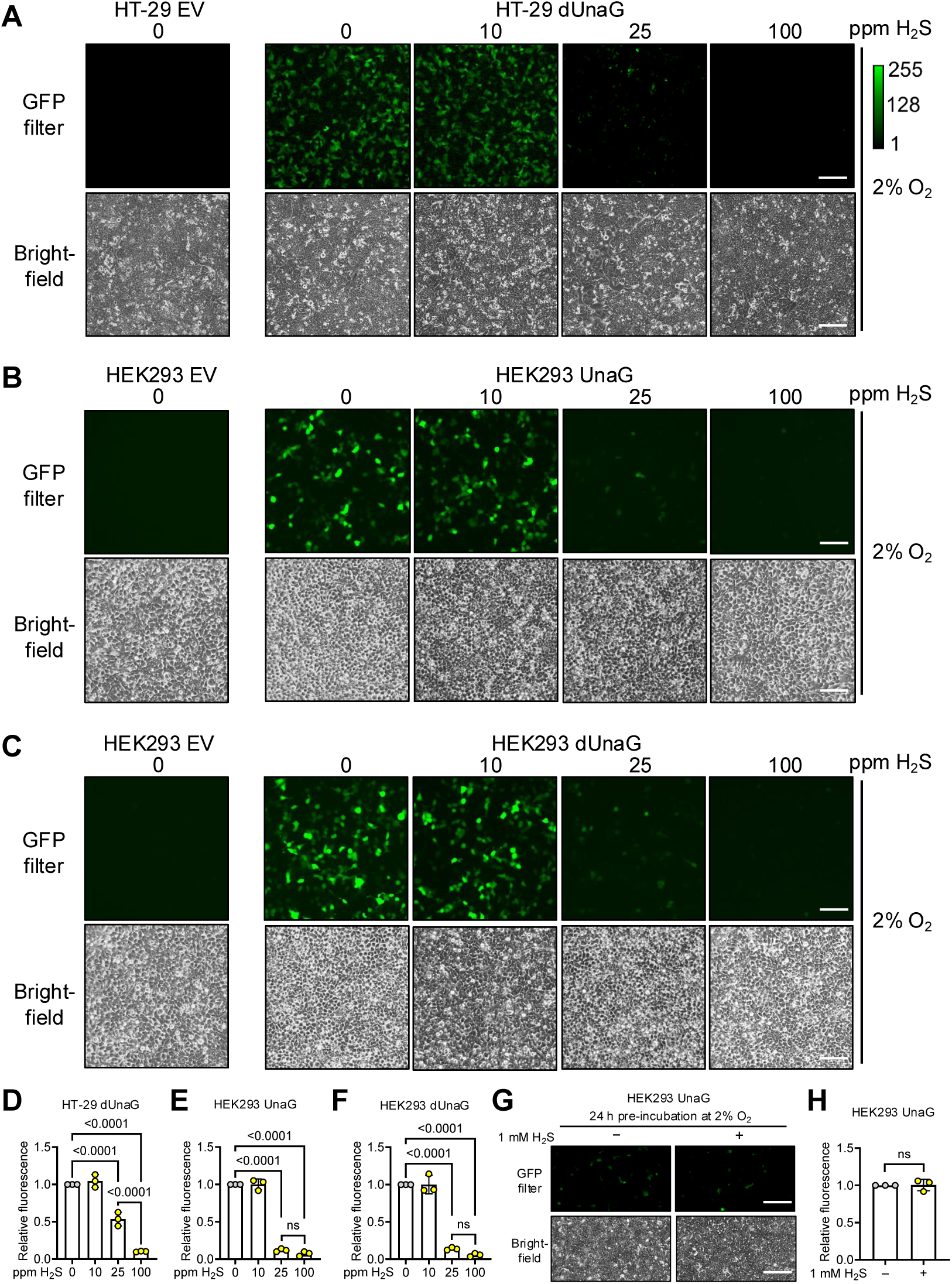
H_2_S-induced increase in O_2_ monitored by dUnaG. (A) Effect of H_2_S (10, 25 and 100 ppm) on dUnaG expression in HT-29 cells grown in 2% O_2_. The images are representative of 3 independent experiments. (B,C) Effect of H_2_S (10, 25 and 100 ppm) on UnaG (B) and dUnaG (C) expression in HEK293 cells grown in 2% O_2_. The images are representative of 3 independent experiments. (D-F) Quantitation of fluorescence intensity in A-C, respectively, normalized relative to the sample lacking H_2_S in each panel. (G,H) Bolus treatment of HEK293 cells expressing UnaG grown in 2% O_2_ with 1 mM H_2_S did not affect fluorescence. Scale bar is 200 μm in all images.

**Supplementary Figure 4:**
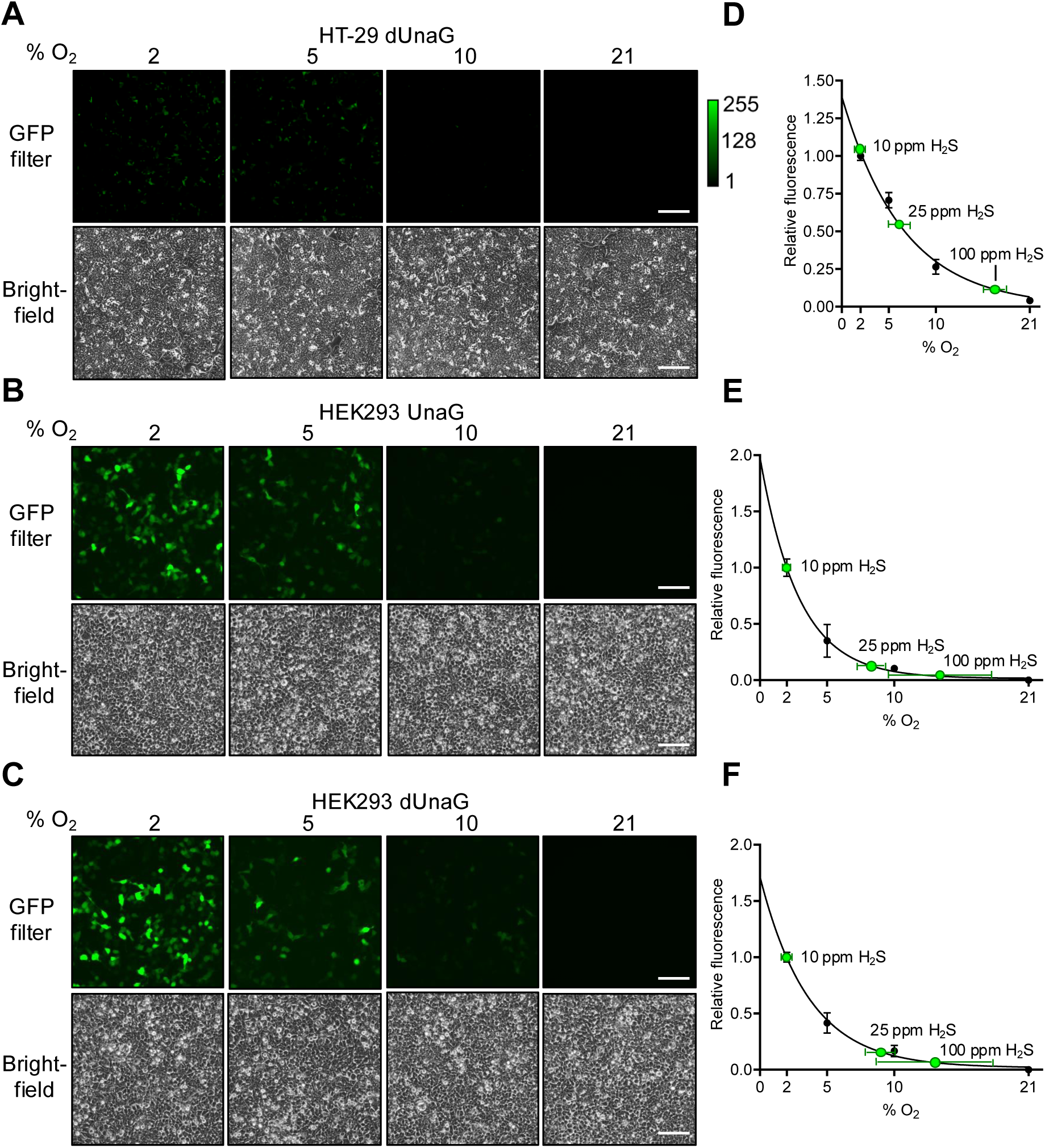
O_2_ calibration curves in HT-29 and HEK cells. (A-C) Expression of dUnaG in HT-29 (A) or HEK293 (C) cells or UnaG in HEK293 cells (B) grown in 2%-21% O_2_ (without H_2_S). (D-F) Dependence of dUnaG (D, F) and UnaG (E) fluorescence on O_2_ concentrations (normalized to 2% O_2_ =1) and fit to an exponential decay curve (n=3). The green dots correspond to fluorescence observed at the indicated concentration of H_2_S (data shown in D-F are from A-C, respectively). Scale bar is 200 μm in all images.

**Supplementary Figure 5:**
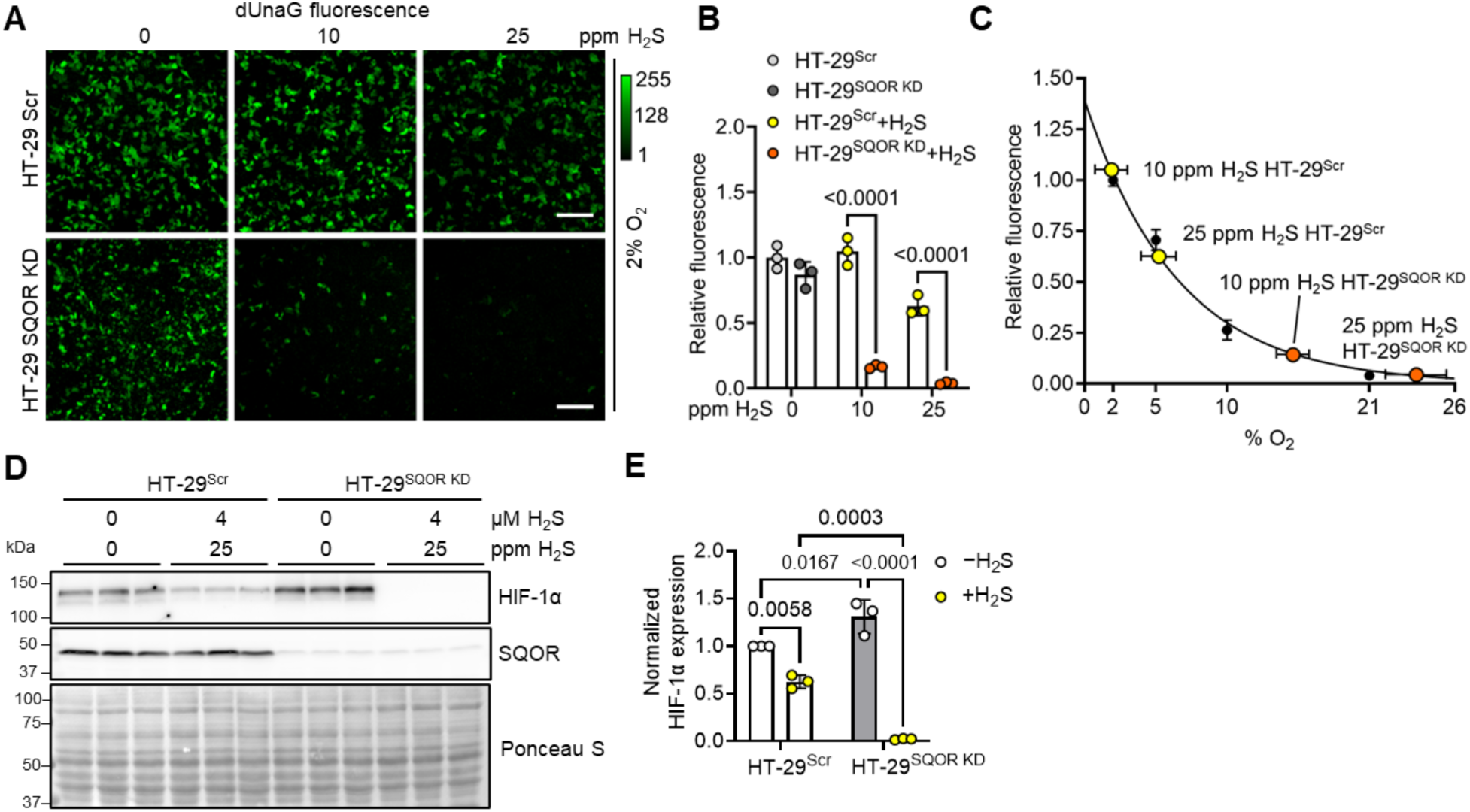
SQOR deficiency sensitizes cells to HIF destabilization by H_2_S. (A) Effect of H_2_S (10 and 25 ppm, 24 h) on dUnaG expression in HT-29^Scr^ and HT-29^SQOR^ ^KD^ cells grown in 2% O_2_. The images are representative of 3 independent experiments. Scale bar is 200 μm. (B) Quantitation of F normalized to HT-29^Scr^ without H_2_S. (C) UnaG fluorescence data in B was plotted on the standard curve for dUnaG fluorescence of O_2_ concentration. (D) HIF-1α and SQOR expression in HT-29^Scr^ and HT-29^SQOR^ ^KD^ cells ± 25 ppm H_2_S exposure for 24 h. (E) Quantitation of HIF-1α levels in D normalized to Ponceau S staining and shown relative to HT-29^Scr^ levels.

**Supplementary Figure 6:**
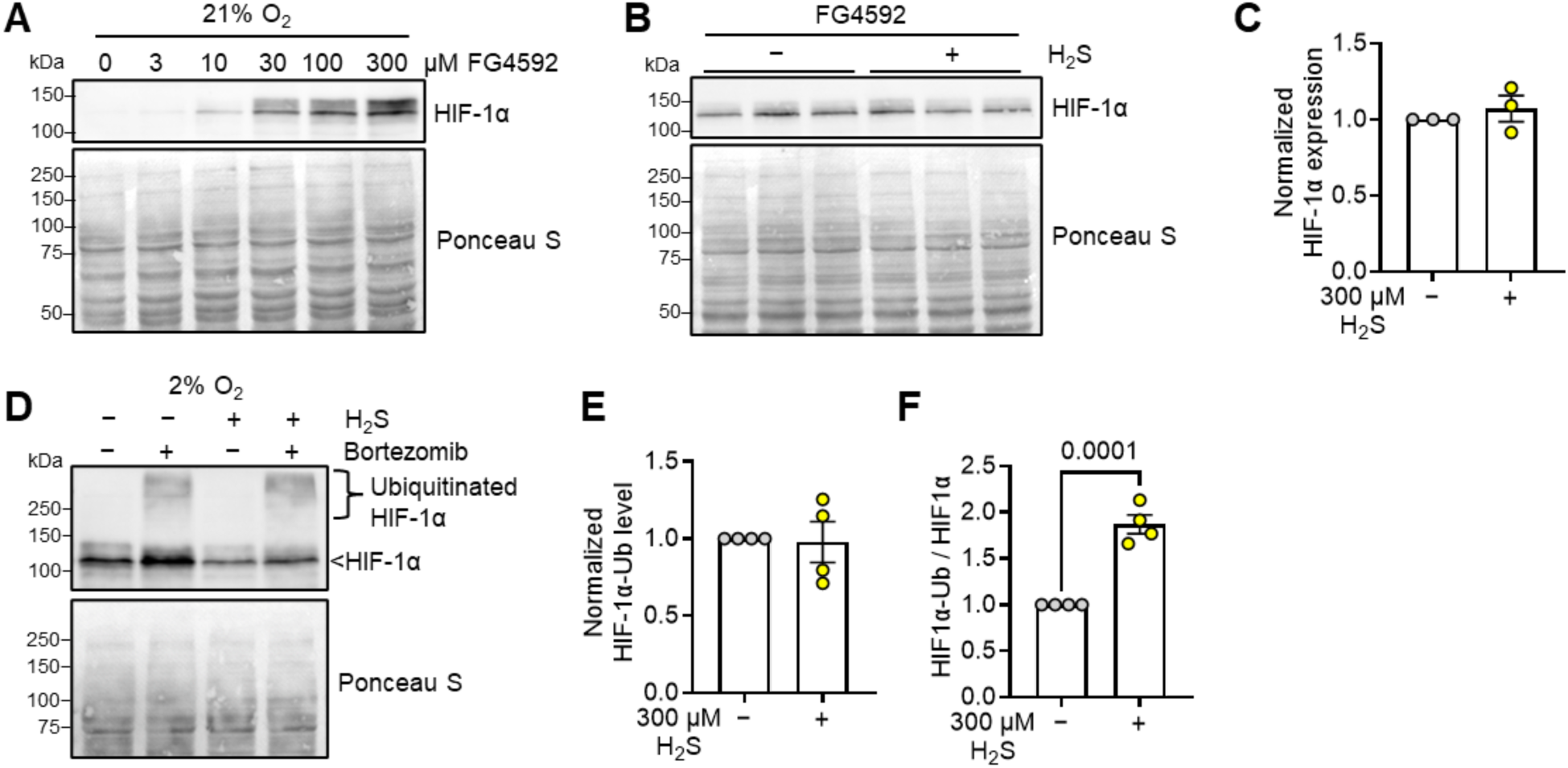
HIF destabilization by H_2_S is PHD-dependent. (A) The PHD inhibitor FG4592 was added to HT-29 cells at the indicated concentrations, and HIF-1α was detected by western blotting. (B) HT-29 cells cultured in 21% O_2_ were treated with 300 µM Na_2_S and 30 µM FG4592, and HIF-1α expression was detected by western blotting. (C) Quantitation of HIF-1α levels in B normalized to Ponceau S staining (n=3). (D) HT-29 cells were pre-treated with the proteasome inhibitor bortezomib (1 µM) for 6 h and then cultured in a 2% O_2_ incubator ± 300 µM Na_2_S for 1 h prior to HIF-1α detection. Accumulation of high molecular weight bands assigned to ubiquitinated HIF was observed in the bortezomib-treated samples. (E,F) Quantitation of HIF-1α levels in D normalized to Ponceau S staining (E) or total HIF-1α levels (F) (n=4).

**Supplementary Figure 7:**
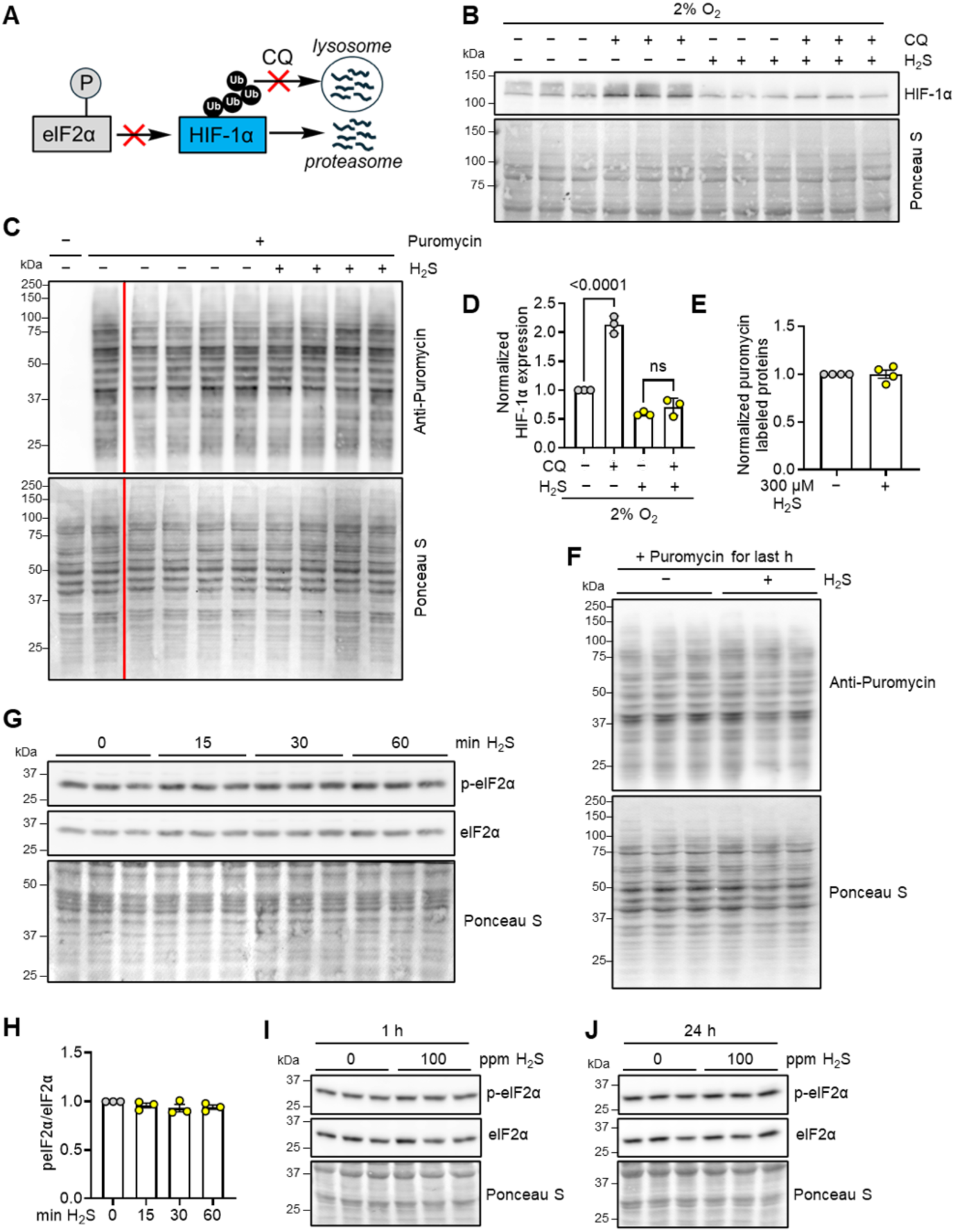
H_2_S does not affect HIF-1α synthesis or its lysosomal degradation. (A) Additional targets for H_2_S-mediated HIF-1α destabilization include decreased translation and lysosomal degradation. (B) HT-29 cells grown in the presence of 2% O_2_ were treated with 50 μM chloroquine (CQ) ± 300 μM H_2_S and HIF-1α was detected by western blot analysis. (C) Quantitation of HIF-1α levels in B normalized to Ponceau S staining (n=3). (D) Puromycin (1 µg/mL, 1 h) led to the accumulation of nascent puromycin-containing peptides, which were not seen in untreated HT-29 cells grown at 21% O_2_. The intensity and banding pattern of the nascent puromycin peptides was unaffected by 300 µM H_2_S. A splice line delineating two separate blots is denoted in red. (E) Quantitation of peptide band intensity in D normalized to Ponceau S staining (n=4). (F) Phosphorylation of eIF2α in HT-29 cells grown in 21% O_2_ and exposed to 300 µM H_2_S for varying times. (G) Quantitation of phospho-eIF2α relative to total eIF2α which was normalized to Ponceau S staining (n=3). (H) HT-29 cells were grown in 21% O_2_ ± 100 ppm H_2_S for 23 h, then incubated for 1 h under the same conditions with 1 μg/mL puromycin. Puromycin-labeled nascent peptides were detected by western blotting. (I,J) Western blot analysis of eIF2α phosphorylation in HT-29 cells grown in the presence of 100 ppm H_2_S and 21% O_2_ after 1 (I) and 24 h (J).

